# How much (ATP) does it cost to build a trypanosome? A theoretical study on the quantity of ATP needed to maintain and duplicate a bloodstream-form *Trypanosoma brucei* cell

**DOI:** 10.1101/2023.03.07.531645

**Authors:** Janaina F. Nascimento, Rodolpho O. O. Souza, Mayke B. Alencar, Sabrina Marsiccobetre, Ana M. Murillo, Flávia S. Damasceno, Richard B. M. M. Girard, Letícia Marchese, Luis A. Luévano-Martinez, Renan W. Achjian, Jurgen R. Haanstra, Paul A. M. Michels, Ariel M. Silber

## Abstract

ATP hydrolysis is required for the synthesis, transport and polymerization of monomers for macromolecules as well as for the assembly of the latter into cellular structures. Other cellular processes not directly related to synthesis of biomass, such as maintenance of membrane potential and cellular shape, also require ATP. The unicellular flagellated parasite *Trypanosoma brucei* has a complex digenetic life cycle. The primary energy source for this parasite in its bloodstream form (BSF) is glucose, which is abundant in the host’s bloodstream. Here, we made a detailed estimation of the energy budget during the BSF cell cycle. As glycolysis is the source of most produced ATP, we calculated that a single parasite produces 6×10^11^ molecules of ATP/cell cycle. Biomass production accounts for ∼62% of the total energy budget, with translation being the most expensive process. Flagellar motility, variant surface glycoprotein recycling, transport and maintenance of transmembrane potential account for less than 30% of the consumed ATP. Finally, there is still ∼9% available in the budget that is being used for other cellular processes of unknown cost. These data put a new perspective on the assumptions about the relative energetic weight of the processes a BSF trypanosome undergoes during its cell cycle.

**Abstract Importance:** Cells use ATP as the main energy currency for the synthesis, organization and maintenance of their macromolecules and cellular structures, in order to stay alive and proliferate. For this purpose, ATP is produced from external nutrients, and is spent by cells in the many processes that are necessary for maintenance and building up new cells. Despite its relevance and the impressive quantity of biological data available, very little is known about how much ATP is required for maintaining and duplicating a cell. In this paper, we present a calculation on how much of the ATP produced by catabolism of the nutrient glucose is used to energize the different processes known to occur during the cell cycle of the infective form of the trypanosomatid parasite that causes human sleeping sickness, the bloodstream form of *Trypanosoma brucei*.

## Introduction

ATP hydrolysis provides most of the free energy used by cells to power biological processes including the metabolic reactions required to build up the biomass for cell proliferation and maintenance. It is possible to estimate the amount of ATP hydrolysis needed for most biological processes and thereby calculate the global ATP expenditure by a cell (Flamholz et al., 2014). During the process of building a new cell, ATP hydrolysis is required for synthesis and polymerization of monomers such as dNTPs and rNTPs for nucleic acids, amino acids for proteins, fatty acids for phospholipids and monosaccharides for oligo- and polysaccharides. ATP hydrolysis is also required for the assembly of complex cell structures such as macromolecular complexes and organelles. Cells may acquire precursors for monomer synthesis or take up ready-to-use monomers from the extracellular environment, but these also require ATP hydrolysis. Furthermore, ATP is necessary for other cellular processes that are not directly related to the synthesis of biomass, such as maintenance of membrane potentials and cellular shape, self-organization, motility, and turnover of molecules.

Parasitic organisms are intriguing in that they may differ in many aspects of their energy expenditure from their free-living counterparts. On the one hand, they may abandon (a sometimes very large) part of their biosynthetic activities if they can acquire multiple nutrients from their host. On the other hand, they may have to invest considerable energy in invasion of the host and in strategies to survive in an environment that tries to tame or kill them (Gadelha et al., 2011). For the present work, we set out to estimate the energy expenditure of the trypanosomatid parasite *Trypanosoma brucei*. *T. brucei* is a unicellular flagellated parasite with a complex life cycle involving insect and mammalian hosts. During its life cycle, *T. brucei* transitions through different cell forms, each one adapted to the specificities of the environment it colonizes. In the gut of the insect vector – the tsetse fly –, amino acids such as proline are abundant and serve preferentially as the energy source for the so-called procyclic trypanosome when glucose is absent (Lamour et al., 2005; Mantilla et al., 2017). In the bloodstream of the mammalian host *T. brucei* can occur in two different developmental forms: long-slender, proliferating trypanosomes and short-stumpy forms. When triggered by a quorum-sensing mechanism, the long-slender trypanosomes differentiate to non-proliferating short-stumpy forms which are competent to develop into procyclic forms when ingested by a tsetse fly (Rojas et al., 2019).

In the blood of the mammalian host, glucose is abundantly available, and it is well established that it is the main source of ATP used by the proliferative bloodstream form (BSF) of the parasite for its proliferation and to survive different environmental challenges (Ryley, 1962; Visser and Opperdoes, 1980). Both procyclic and bloodstream forms of *T. brucei* can be easily cultivated *in vitro* in semi- or completely defined media (Creek et al., 2013; Hirumi and Hirumi, 1989), which has enabled the detailed investigation of the end-products obtained from different substrates as well as the estimation of metabolic fluxes. In these organisms, the major part of the glycolytic pathway is compartmentalized in peroxisome-related organelles called glycosomes (Michels et al., 2021; Opperdoes and Borst, 1977). Noteworthy, while procyclic forms can oxidize metabolites (including glucose-derived pyruvate) in their single mitochondrion, under most conditions the BSF obtain their energy by aerobic fermentation with no involvement of oxidative phosphorylation.

The total energy cost of a biological process can be expressed as the summation of the direct costs (amount of the necessary ATP hydrolysis) spent on all energy-requiring processes (Mahmoudabadi et al., 2019). In contrast to most bacteria and yeasts, BSF *T. brucei* use very little of the glucose consumed to synthesize biomass (Haanstra et al., 2012). Noteworthy, these trypanosomes depend on extracellular availability of other essential nutrients to serve as carbon sources for the biosynthesis of precursors of macromolecules for biomass. Thus, knowledge of the rate of glucose consumption, together with the fact that almost all glucose consumed by the BSF is directed to ATP formation allows calculation of the total amount of ATP produced per cell cycle. We can also estimate the ATP expenditure during a cell cycle as other relevant parameters are known such as doubling time, molecular content, genome size, transcriptome and proteome half-lives, and cell motility.

For some free-living prokaryotic and eukaryotic microorganisms calculations of metabolic energy obtained (mostly transduced into ATP) from external sources have been reported previously (Lynch and Marinov, 2017, 2015; Stouthamer, 1973).These calculations included energy obtained from external sources (oxidation of organic or inorganic molecules; absorbance of light) through different processes and the energy used for different activities (biosynthesis of macromolecules, biogenesis of (sub)cellular structures, transmembrane transport of molecules, motility, *etc*.). Here, we present a detailed estimation of the energy (ATP) budget and the energy costs of the two main commitments that a long-slender BSF *T. brucei* has during a cell cycle: to stay alive (maintenance) and to make a new cell (duplication). We found that the production of biomass, including the turnover of parts of its components under standard cultivation conditions, accounts for approximately 62% of the energy budget, with translation being the most “expensive” process. We estimated the extent to which several other cellular processes are responsible for using the remaining ATP that these cells produce.

## Results

### How much ATP is produced by *T. brucei* BSF during a cell cycle?

The BSF *T. brucei* model studied

The BSF of *T. brucei* is one of the relevant trypanosomatids for public health, and the availability of data about the various activities it exerts when parasitizing its mammalian hosts, such as proliferation, catabolic and anabolic processes, endocytosis, motility, among others led us to select it to estimate its ATP budget for cell maintenance during a cell cycle and for making an entirely new cell. Most data used for calculation of the ATP production have previously been obtained by using *T. brucei* strain Lister 427, BSF cell line 449 (Haanstra et al., 2012). Trypanosomes of this Lister 427 strain are monomorphic, with the BSF occurring only as proliferating long-slender forms because they are incapable of differentiating to stumpy forms. Within specific cell densities *in vitro* growth is exponential and the specific glycolytic flux is constant (Haanstra et al., 2012). For the costs of making the building blocks of the cell such as dNTPs and amino acids, we used available data on the characterized biosynthetic pathways as well as the genome annotation for the presence of still uncharacterized pathways. For those biological processes in which energy costs are not yet fully understood for *T. brucei*, we made inferences based on data available for other organisms.

As previously mentioned, BSF *T. brucei* rely (almost) completely on glycolysis for their energy requirements and excrete nearly all pyruvate produced rather than further oxidizing it in the mitochondrion (Haanstra et al., 2012). The first seven enzymes of the glycolytic pathway are compartmentalized in peroxisome-related organelles called glycosomes (Opperdoes and Borst, 1977). The reoxidation of the glycolytically produced NADH occurs through the transfer of the electrons by a shuttle mechanism from the glycosomes to the mitochondrion, in which glycolytically produced dihydroxyacetone phosphate is reduced to glycerol 3-phosphate with the concomitant oxidation of NADH to NAD^+^ by a glycosomal glycerol-3-phosphate dehydrogenase. In turn, the produced glycerol-3-phosphate is oxidized back to dihydroxyacetone phosphate by a mitochondrial glycerol-3-phosphate dehydrogenase, with the concomitant reduction of FAD to FADH_2_ which, in aerobic conditions, is reoxidized to FAD by the transfer of electrons to oxygen catalyzed by the trypanosome alternative oxidase (Helfert et al., 2001). Summarizing, this shuttle occurs without classical oxidative phosphorylation (OxPhos) (Michels et al., 2021; Opperdoes and Borst, 1977). In fact, in this stage of the parasite’s life cycle, enzymes of the tricarboxylic acid (TCA) cycle are either absent or severely downregulated (Zíková et al., 2017), and the F_1_F_O_-ATP synthase complex works in “reverse mode” accounting for an H^+^/ATPase activity pumping protons into the intermembrane space, for the maintenance of the mitochondrial membrane potential (Nolan and Voorheis, 1992; Schnaufer et al., 2005). Due to the absence of classical OxPhos, glycolysis is the main source of ATP in BSFs (Opperdoes, 1987). Net production of ATP, and thus the free-energy yield of glycolysis occurs in the cytosol and almost entirely comes from the flux through the enzyme pyruvate kinase (Haanstra et al., 2012). It has been shown that some ATP synthesis can occur in the mitochondrion by the acetate:succinate CoA transferase / succinyl-CoA synthetase (ASCT/SCS) cycle, which can use as a substrate acetyl-CoA derived from relatively minute amounts of pyruvate routed to the mitochondrion and/or from threonine oxidation. However, the amount of ATP produced by this system is small when compared to that produced by glycolysis and may vary depending on conditions. (Michels et al., 2021; Mochizuki et al., 2020). Taking all this information into account, we can proceed to make a reliable estimation of the total amount of ATP that is produced during a complete cell cycle, in which an entire *Trypanosoma* cell is built.

According to data from Haanstra et al. (2012) when BSF *T. brucei* strain Lister 427, cell line 449 was growing exponentially in HMI-9 medium (for composition see Supplementary Table S1) at 37 °C in the presence of 25 mM of glucose, the glucose consumption flux was 160 nmol/(min × 10^8^ cells). As mentioned, virtually all consumed glucose (155.2 nmol/min × 10^8^ cells) was directed towards pyruvate under aerobic conditions. However, it should be noted that, depending on the culture conditions, a small part of glycolytically-derived metabolites can be used for the synthesis of sugar nucleotides (Turnock and Ferguson, 2007), inositol (Martin and Smith, 2006), acetate (Creek et al., 2015; Mazet et al., 2013), amino acids such as asparagine and alanine (Creek et al., 2015), which can contribute to anabolic processes. Stoichiometrically, the glycolytic breakdown of one molecule of glucose yields two molecules of pyruvate, and each of these is accompanied with the yield of one ATP, resulting in an ATP synthesis flux of 310.4 nmol/(min × 10^8^ cells). This flux remains constant throughout the exponential proliferation phase (Haanstra et al., 2012), and therefore we calculated the total amount of ATP produced by one cell during one cell cycle (5.3 h in the experiment by Haanstra et al. 2012; for details see Materials and Methods), which results in 6.00 × 10^11^ molecules of ATP/(cell cycle * cell). It is worth remarking that, despite the concentration of glucose in this culture medium is high with respect to that present in the mammalian blood, the BSF proliferation in this culture condition can be compared with the BSF proliferation in blood, since both, the medium and the blood glucose concentration (∼5 mM), are more than enough to saturate the glucose uptake in these cells (Eisenthal et al., 1989; Jean Gruenberg et al., 1978; ter Kuile and Opperdoes, 1991a; Tetaud et al., 1997). The fact that the population doubling time described by Haanstra et al. is very similar to that reported previously for different *T. brucei* strains including Lister 427 in mammalian blood supports the relevance of these data (Michels, unpublished results, Doyle et al., 1980; Miller and Turner, 1981).

### The cost of genome duplication

To express and transmit its genetic information, every cell needs to duplicate and spatially organize its DNA, transcribe the information into RNA, and translate it into functional proteins. The energy requirements of each of these processes differ and include the costs of making, assembling, and processing the building blocks of each polymer. Cells duplicate their genome once during the cell cycle, which requires activated nucleotides. It has been established for yeast and bacteria that the cost of synthesis of all requisite nucleotides *de novo* from glucose is approximately 50 ATPs per nucleotide (Lynch and Marinov, 2015). Trypanosomatids lack the purine *de novo* biosynthetic pathway (Berens et al., 1981) and therefore rely on the purine salvage pathway by import of appropriate nitrogenous bases to be used as precursors for the synthesis (Davies et al., 1983). In addition, trypanosomatids can synthesize pyrimidines from glutamine and aspartate, both present in the culture medium HMI-9. So far, there is no evidence that it can import thymidine or thymine (reviewed in Tiwari and Dubey, 2018). Based on the metabolic pathways predicted from the *T. brucei* genome for purine salvage and pyrimidines biosynthesis we calculated the ATP cost for the biosynthesis of each nucleotide (Table 1), starting from the precursors available in the culture medium: hypoxanthine (for purine salvage) and glutamine and aspartate (for the *de novo* synthesis of pyrimidines). The direct costs of making the other metabolites required in these pathways were also included (Supplementary Tables S2, S3 and S4). On average, *T. brucei* spends 11.5 ATP molecules for the biosynthesis of one purine and 9 ATPs for the biosynthesis of one pyrimidine (Table 1). The *T. brucei* haploid genome has an approximate size of 35 Mbp (TriTrypDB; https://tritrypdb.org/tritrypdb/app) and consists of 11 megabase chromosomes, a few intermediate chromosomes, and hundreds of minichromosomes (Berriman et al., 2005). Given the cost of each dNTP and the GC content of the *T. brucei* genome, the estimated total cost of the synthesis of the necessary number of dNTPs for the entire diploid genome duplication in one cell cycle is then 1.4 × 10^9^ ATPs.

**Table 1.** ATP cost for the synthesis of deoxyribonucleotides for *T. brucei* genome duplication.

| dNTP | ATP cost | % of the genome | Total cost |
| --- | --- | --- | --- |
| dCTP | 12 | 22.8 | $3.8 \times 10^8$ |
| dTTP | 6 | 27.2 | $2.3 \times 10^8$ |
| dATP | 11 | 27.2 | $4.2 \times 10^8$ |
| dGTP | 12 | 22.8 | $3.8 \times 10^8$ |
| **Total** |  |  | **$1.4 \times 10^9$** |

Other costs involved in genome duplication were estimated. First, there is the cost of the unwinding of the double-helix of the DNA. Using the yeast value, where this process costs one ATP per nucleotide (Ramanagoudr-Bhojappa et al., 2013), in *T. brucei* it will require 7 × 10^7^ ATPs in total. Next, some ATP is needed for the synthesis of the small RNA primers (∼10 nt) necessary for the initiation of nucleotide polymerization during duplication of the lagging strand of DNA, which involves the formation of the Okasaki fragments. The number of the necessary RNA primers depends on the number of the origins of replication (ORI) and the size of the intervals between them. In yeast, the length of the Okasaki fragment is ∼165 nt, with 10 nt corresponding to the RNA primer (Smith and Whitehouse, 2012). Taking that: i. the haploid genome has 35 Mb; ii. the lagging strand during DNA replication is fully replicated based on the synthesis of Okasaki fragments; and iii. that each Okasaki fragment has a length of ∼165 nt, the total number of Okasaki fragments needed for the genome replication can be obtained from the ratio between the genome size and the length of the Okasaki fragment. The obtained value indicates that 4.2 × 10^5^ is the minimum number of RNA primers necessary to produce the Okasaki fragments necessary to duplicate the whole diploid genome. The average cost of rNTP synthesis in *T. brucei* is 5 ATPs per unit (see below). Therefore, the costs associated with RNA primer synthesis are 2.1 × 10^7^ ATPs. After the synthesis of Okasaki fragments, DNA ligase uses 2 ATPs to ligate each pair of fragments, which then costs 8.4 × 10^5^ ATPs in this parasite. Last, there is an ATP cost associated with the assembly of the polymerase-containing sliding clamp. On average, 3 ATPs per complex are necessary (Majka et al., 2004). Since duplication of the lagging strand requires one sliding clamp per fragment to be synthesized, *T. brucei* requires approximately 1.3 × 10^6^ ATPs in this step. As a whole, the contribution of these processes to the total cost is minor when compared to the cost of nucleotide synthesis (Table 2).

**Table 2.** Summary of ATP costs associated with nuclear and mitochondrial genome duplication (maxicircles and minicircles) of *T. brucei*.

| Process | ATP cost |  |  |
| --- | --- | --- | --- |
|  | Nuclear genome | Maxicircles | Minicircles |
| dNTP synthesis | $1,400 \times 10^6$ | $6.9 \times 10^6$ | $60 \times 10^6$ |
| DNA unwinding | $70 \times 10^6$ | $0.69 \times 10^6$ | $6 \times 10^6$ |
| RNA primer synthesis | $21 \times 10^6$ | $0.21 \times 10^6$ | $1.8 \times 10^6$ |
| Okasaki fragments ligation | $0.84 \times 10^6$ | $0.0084 \times 10^6$ | $0.073 \times 10^6$ |
| Sliding clamp assembly | $1.3 \times 10^6$ | $0.012 \times 10^6$ | $0.11 \times 10^6$ |
| Opening of ORIs | negligible | negligible | $0.12 \times 10^6$ |
| Total | **$1,493 \times 10^6$** | **$7.8 \times 10^6$** | **$68.1 \times 10^6$** |

There is still a series of costs that is too small to be relevant to the total cost of genome duplication. One example is the ATP investment associated with opening the ORIs. It has been estimated as being at least 20 ATPs per ORI (Lynch and Marinov, 2015). In *T. brucei*, there is a minimum number of 33 ORIs necessary to replicate the 11 megabase chromosomes (da Silva et al., 2020), which adds at least 1,320 ATP molecules per S-phase of the cell cycle. Additionally*, T. brucei* has at least 6 intermediate-sized chromosomes and about 50-100 minichromosomes (Melville et al., 2000). Assuming that there is at least one ORI per intermediate and minichromosome, there will be an additional requirement of about 620 to 2,120 ATP molecules. Other costs such as for proofreading, DNA repair, and epigenetic modifications are still to be fully elucidated. The total cost for the nuclear genome duplication is estimated as being 1.49 × 10^9^ ATP molecules.

### The cost of kDNA duplication

The mitochondrial genome of *T. brucei* is contained in a unique structure called kinetoplast. The DNA present in the kinetoplast (kDNA) consists of a concatenated network of two classes of circular DNA: the maxicircles (∼23 kb) and minicircles (∼1 kb). Maxicircles are present in a low-copy number (∼30 per cell) and encode proteins of the mitoribosomes, some of the proteins of the complexes of the respiratory chain, and two rRNAs. Remarkably, most of these genes in maxicircles are encrypted and need to undergo RNA editing before translation. The RNA editing process is mediated by guide RNAs (gRNAs) that are transcribed from the minicircles. There are approximately 6,000 minicircles per cell with at least 391 different sequences encoding different gRNAs (Cooper et al., 2019). Because of the intricate nature of the kDNA, the process of its duplication is rather complex. On one hand, minicircles are released from the core of the network, unwound, duplicated and then reassembled back in the periphery of the network. On the other hand, maxicircles are duplicated inside the network, but the exact mechanism is still unknown (reviewed in Verner et al., 2015).

As has been described for genome replication, dozens of proteins participate in kDNA duplication, including helicases, topoisomerases, polymerases, primases and ligases (reviewed in Jensen and Englund, 2012). As the same classes of proteins are involved in both processes, we assumed similar costs for the initiation of each replication unit to those estimated for the nuclear genome duplication. Therefore, we used the rationale and estimations described in the previous section: (i) DNA unwinding, which costs 1 ATP per nucleotide, resulting in 0.69 × 10^6^ ATPs for maxi- and 6 × 10^6^ ATPs for minicircle duplication; (ii) RNA primer synthesis costs 50 ATPs per primer, resulting in 0.21 × 10^6^ ATPs for maxi- and 1.8 × 10^6^ ATPs for minicircles; (iii) Okasaki fragments ligation costs 2 ATP per ligation resulting in 0.0084 × 10^6^ ATPs for maxi- and 0.073 × 10^6^ ATPs for minicircles; and (iv) sliding clamp assembly which costs 3 ATPs on average, resulting in 0.012 × 10^6^ ATPs for maxi- and 0.11 × 10^6^ ATPs for minicircles (Table 2).

Some peculiarities regarding the kDNA and its replication required an adjustment in the calculations. First, although the sequence of kDNA is mostly known, the distribution of the 391 types of minicircles varies from 1 to 144 copies per cell (Cooper et al., 2019). This makes the accurate GC-content hard to estimate. For this reason, we assumed a 50% CG content and an average synthesis cost of 10 ATPs per nucleotide. Thus, the cost of the dNTPs for maxicircle duplication is 6.9 × 10^6^ ATPs and 60 × 10^6^ ATPs for minicircle duplication. Second, according to the calculations made for nuclear genome DNA replication, the cost for opening the origins of replication is 20 ATPs per ORI. We have previously considered this cost negligible due to the low number of ORIs necessary to duplicate the whole nuclear genome. Although this number is still negligible for the duplication of maxicircles (∼600 ATPs), due to the number of minicircles (20 ATPs per ORI for 6,000 ORIs), this cost becomes more relevant for their duplication, and it totalizes 0.12 × 10^6^ ATP molecules (Table 2). The duplication of the mitochondrial genome (maxicircles and minicircles) costs 0.0759 × 10^9^ATP molecules. In total, duplicating both the nuclear and the mitochondrial genome requires an estimated 1.57 × 10^9^ ATP molecules.

### The cost of transcription of the nuclear genome

In *T. brucei* BSF, RNA Pol I transcribes the gene arrays for ribosomal RNAs (rRNAs) and a telomeric expression site containing a single variant surface glycoprotein (VSGs) gene. This specific gene comes out of a very large repertoire of which one VSG is expressed at a given time. However, together with this VSG gene, a set of genes called Expression Site Associated Genes (ESAGs) are transcribed that lie upstream of the VSG gene (Bernards et al., 1985; Johnson et al., 1987; Kooter et al., 1987). Most of them encode proteins with still unknown biological function. RNA Pol II transcribes all other protein-coding genes as well as the genes for a spliced leader (SL) RNA, whilst RNA Pol III transcribes genes encoding snRNAs, tRNAs, and 5S RNAs (Gilinger and Bellofatto, 2001; Günzl et al., 2003). In trypanosomatids, genes are organized in tandem arrays which are transcribed in a polycistronic manner. The resulting long precursor RNAs are processed by trans-splicing and polyadenylation. Consequently, mature individual mRNAs containing a 39 nt SL with a 5’cap and a 3’poly-A tail are produced (Jäger et al., 2007). It means that, differently from organisms that regulate transcription initiation and termination of each gene, trypanosomatids transcribe coding genes that are not needed in a specific condition (*e.g.* the tandemly arranged genes encoding PGKA, B and C are all transcribed simultaneously, but B or C is degraded depending on the life-cycle stage (Gibson et al., 1988; Haanstra et al., 2008; Osinga et al., 1985), as well as intergenic regions and then degrade them once the mature mRNAs are formed.

Regarding the ATP costs of transcription, we estimated the ATP costs for synthesis of an entire set of transcripts and the ATP costs associated with their maintenance (turnover). For that purpose, we used most of the data and assumptions used for the model developed by Haanstra and collaborators for different aspects of BSF *T. brucei* gene expression (Haanstra et al., 2008). In this paper they also report values and estimations for numbers and half-lives of four types of RNAs: i. rRNAs; ii. RNAs encoding VSGs; iii. mRNAs; iv. SL-RNAs (Table 3). For the ATP expenditure calculation, we considered the cost of synthesis of the rNTPs to be used as monomers, the cost of each polymerization reaction, and the steady-state number of molecules of each RNA-type produced per cell (*N*) and the average length of the mature RNA (*L*).

**Table 3.**
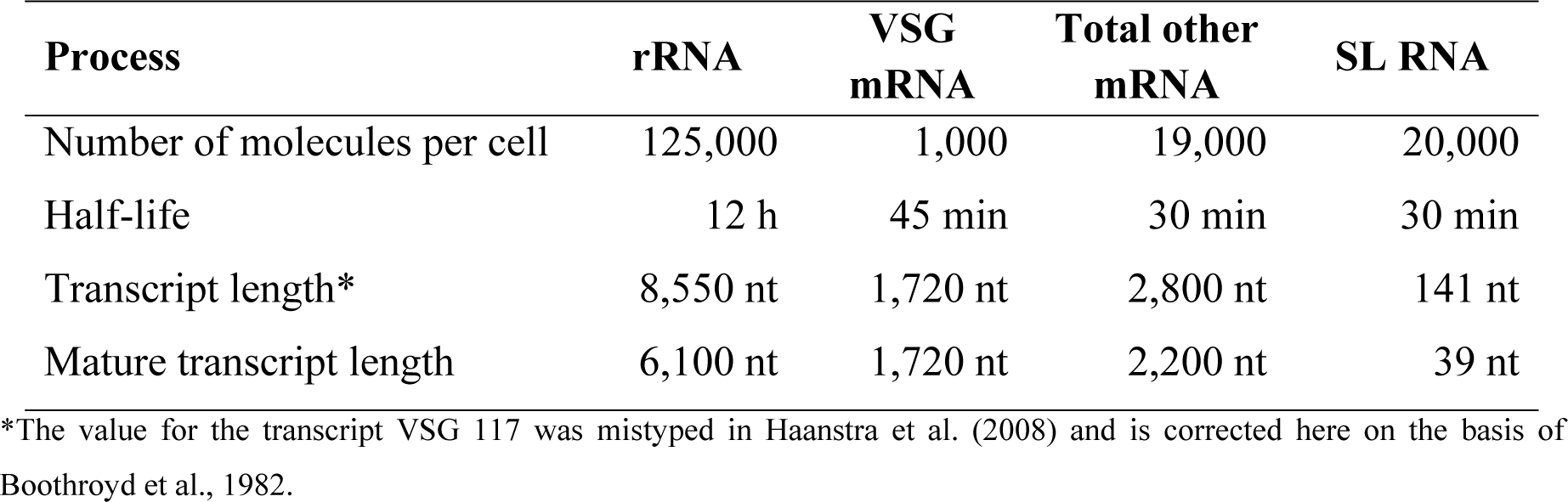
Data from Haanstra et al., 2008 used in this work.

| Process | rRNA | VSG mRNA | Total other mRNA | SL RNA |
| --- | --- | --- | --- | --- |
| Number of molecules per cell | 125,000 | 1,000 | 19,000 | 20,000 |
| Half-life | 12 h | 45 min | 30 min | 30 min |
| Transcript length* | 8,550 nt | 1,720 nt | 2,800 nt | 141 nt |
| Mature transcript length | 6,100 nt | 1,720 nt | 2,200 nt | 39 nt |

\*The value for the transcript VSG 117 was mistyped in Haanstra et al. (2008) and is corrected here on the basis of Boothroyd et al., 1982.

#### Synthesis of the transcriptome

The production cost of the nucleotides is on average 5 ATPs per rNTP (Table 4, Supplementary Table S3). The total synthesis cost for the four RNA populations is 5*NL*. Therefore, the resulting ATP cost for the (Lynch and Marinov, 2015) rRNA population is 3.8 x10^9^, for the VSG mRNAs 8.6 x10^6^, for the set of other mRNAs 2.1 x10^8^ and for SL RNA synthesis 3.5 × 10^6^ per cell cycle (Table 5).

**Table 4.** ATP cost for the synthesis of ribonucleotides.

| rNTP | ATP cost |
| --- | --- |
| CTP | 5 |
| UTP | 4 |
| ATP | 5 |
| GTP | 6 |

**Table 5.**
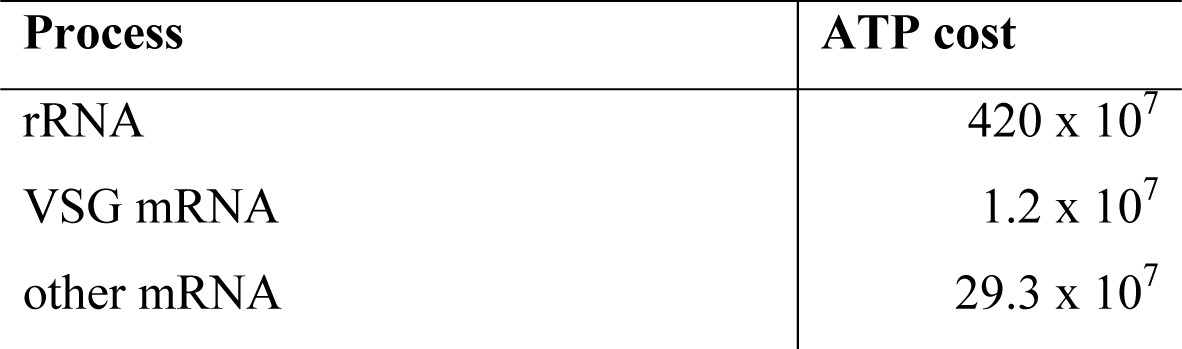
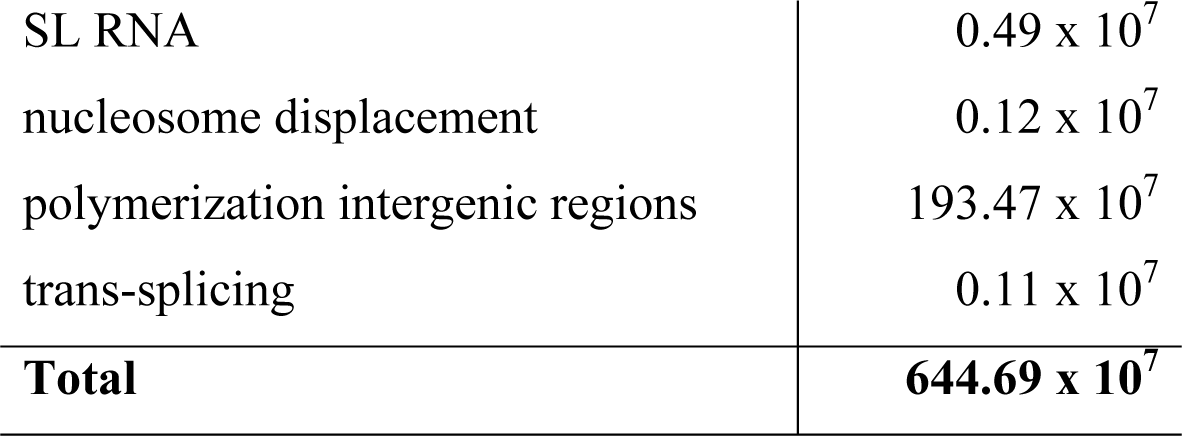
Summary of ATP costs associated with transcription per cell cycle of *T. brucei*.

| Process | ATP cost |
| --- | --- |
| rRNA | $420 \times 10^7$ |
| VSG mRNA | $1.2 \times 10^7$ |
| other mRNA | $29.3 \times 10^7$ |
| SL RNA | $0.49 \times 10^7$ |
| nucleosome displacement | $0.12 \times 10^7$ |
| polymerization intergenic regions | $193.47 \times 10^7$ |
| trans-splicing | $0.11 \times 10^7$ |
| **Total** | **$644.69 \times 10^7$** |

#### Maintenance of the transcriptome (turnover)

Assuming that ribonucleotides are efficiently recycled, the cost invested in recharging the rRMPs to rRTPs is 2 ATPs (Lynch and Marinov, 2015). Considering the half-lives (t_1/2_) of each set of RNAs, the maintenance cost is the cost of replacing the RNAs degraded during the cell cycle. Given the doubling time of BSF *T. brucei*, here taken as 5.3 hours (see above) and the half-life of each set of RNA, we calculated the number of RNA molecules of each class that must be resynthesized during a cell cycle for replacement (N_r_) as being 32,965 for rRNAs, 993 for VSG mRNAs, 18,988 for every other mRNA and 19,987 for the SLs. Hence, as we considered a complete recycling of the ribonucleotides obtained from the RNA degradation (NMPs), the cost for maintaining the whole transcriptome is the cost of recharging the nucleotides to be polymerized. For each RNA subset we calculated the cost as 2*N_r_L*. According to this, the total cost for the maintenance of each type of RNA is 40 × 10^7^ for rRNAs and 0.34 × 10^7^ ATPs for VSG mRNAs, whereas the maintenance of the remaining set of mRNAs costs 8.3 × 10^7^ ATPs, and the cost calculated for SL-RNA is 0.14 × 10^7^ ATPs (Table 5).

#### Polymerization of rNTPs of intergenic regions

As the intergenic regions are transcribed and degraded to monomers after RNA processing, we considered that the ribonucleotides used in the transcription of intergenic regions are efficiently recycled. However, the cost invested in polymerizing the ribonucleotides of the intergenic regions must be estimated. For this purpose, we used the difference in length between the whole precursor and the mature transcripts and applied the same calculations for synthesis and maintenance for the polymerization costs (Haanstra et al. 2008). For the VSG transcripts, the whole transcript length is considered as being the same of the mature transcript length (Boothroyd et al., 1982; Haanstra et al., 2008). However, VSG genes are transcribed together with the ESAGs in a polycistronic manner in one out of the about 15 telomeric bloodstream expression sites (BES) that is activated. Therefore, we calculated the total length of the intergenic regions of the polycistron. For this, we used data from the BES 40 containing the VSG 221 gene (Müller et al., 2018). The whole BEŚs length is 59.78 kb. It contains 18 protein-coding sequences including the VSG with a total added size of 25.15 kb. For the estimation of the UTR regions (which also constitute the mature RNA) we used the median length of 130 nt for the 5’UTR and 399 nt for the 3’ UTR (Michaeli, 2011) except for the VSG, where we considered the whole size of 1,720 nt (Boothroyd et al., 1982; Haanstra et al., 2008). So, the total length of the polycistron that is maintained as mRNA is 34.44 kb. Therefore, the intergenic regions that are transcribed and further degraded are estimated as being 25.34 kb long. Applying the same calculations for synthesis and maintenance for the polymerization used above, we estimated the total cost of intergenic transcription for VSG/ESAGs, rRNA, mRNAs of other proteins, and SL RNA as being 18 × 10^7^, 166 × 10^7^, 8 × 10^7^ and 1.47 × 10^7^ ATPs, respectively, per cell cycle (Table 5).

#### Nucleosome displacement

Another cost associated with transcription is related to the displacement of the nucleosomes. This process involves various histone posttranslational modifications (Stillman, 2018). *T. brucei* expresses four out of five canonical eukaryotic variants of histones (H2A, H2B, H3, and H4) and they work as boundaries for polycistronic units (Lowell et al., 2005; Siegel et al., 2009). The length of DNA wrapped around each nucleosome is ∼147 nt and the length of the strands linking two nucleosomes is ∼43 nt in *T. brucei* (Hecker et al., 1989). Considering these numbers and the total DNA length, the number of nucleosomes can be estimated as being 3.7 × 10^5^ per diploid genome. Assuming a minimum cost of 30 ATPs per set of modifications in one nucleosome (Lynch and Marinov, 2015) and that once the chromatin is open for transcription it remains in this state, the minimum cost of displacing the nucleosome barriers during transcription is 1.1 × 10^6^ ATPs per cell cycle (Table 5).

*Splicing:* By far the major part of the mRNA maturing process occurs by trans-splicing (with only two exceptions reported (Siegel et al., 2010)). In trans-splicing, similarly to cis-splicing, two transesterification reactions unite two RNA fragments (reviewed in Michaeli, 2011). Cis-splicing costs at least 10 ATPs per intron (Lynch and Marinov, 2015; Matera and Wang, 2014) and here we consider the same cost for trans-splicing. Considering the synthesis and maintenance of mRNA levels, the cost of trans-splicing is 1.06 × 10^6^ ATPs per cell cycle (Table 5).

In summary, transcription costs ∼1.1 × 10^10^ ATP molecules. Costs associated with other aspects of transcription such as the formation of the transcriptional complexes are too small or have not been completely elucidated and therefore are not considered here. In eukaryotes, RNA polymerase II transcription initiates with the recruitment of the polymerase to the promoter region by multiple transcription factors. Subsequently, the DNA helix is unwound, forming an open complex (OC). These processes cost at least 20 ATPs per OC (Lynch and Marinov, 2015; Wang et al., 1992; Yan and Gralla, 1997). Because of the polycistronic transcription, fewer OCs are necessary to initiate transcription in trypanosomatids, making these costs negligible to the total transcriptional cost. Similarly, transcriptional termination is likely to be less costly in trypanosomatids, since it happens at transcription termination sites marked by histone variant H3.V and base J, a modified thymine detected in the nuclear DNA of trypanosomatids and related protists grouped in the Euglenozoa clade (Reynolds et al., 2016; Schulz et al., 2016). Additionally, some transcriptional costs have not been completely elucidated. For example, phosphorylation of the C-terminal domain of RNA Pol II regulates different aspects of transcription (Hsin and Manley, 2012). However, the number of phosphorylation events per transcriptional cycle in trypanosomatids has not been determined yet. Another process related to transcription of which exact costs remain unknown is RNA nuclear export. Interestingly, although this process is ATP dependent in opisthokonts (Folkmann et al., 2011), the lack of many ATPases in the nuclear pore complex of trypanosomatids suggests that mRNA nuclear export is GTP driven in these organisms (Obado et al., 2016). Regardless of the case, these costs remain to be determined.

### The costs of transcription of kDNA

The maxicircles of the kDNA code for 2 rRNAs and 18 proteins (Kirby et al., 2016). It is currently accepted that, similarly to what happens in the trypanosomatid nucleus and mitochondria of other organisms, transcription of the maxicircles is polycistronic and that the long pre-RNAs are processed at both ends to generate mature RNAs (Gazestani et al., 2018; Koslowsky and Yahampath, 1997). However, it has been recently proposed that this transcription might be gene-specific and promoter-regulated (Aphasizheva et al., 2020; Sement et al., 2018). Additionally, 12 of these genes, named cryptogenes, need to undergo further processing by RNA editing to generate translation-competent mRNAs. This editing consists of the insertion and/or deletion of uridines and is mediated by gRNAs transcribed from the minicircles present in the kDNA (reviewed in Read et al., 2016). Once transcribed, these gRNAs are also processed by 3’-5’ trimming and U-tailing and stabilized by their ligation to the RNA-editing substrate-binding complex (RESC) (reviewed in Aphasizheva et al., 2020). Multiple gRNAs are necessary for the editing of a single maxicircle-encoded mRNA (Koslowsky et al., 2014).

To estimate the minimal cost of kDNA transcription, and due to the lack of data on the number of kDNA transcripts per BSF cell and their half-lives, we assumed that maxicircle transcription has similar dynamics to that of nuclear transcription. Noteworthy, most of the mtDNA genes are developmentally regulated but, in the model of polycistronic transcription, this regulation is likely to be posttranscriptional (Gazestani et al., 2018). Thus, considering a similar ratio of the number of transcripts/genes to the nucleus, and the number of maxicircles (∼30) present in the kDNA, we estimated an average of 480 molecules of mRNA and 35,700 molecules of rRNA per BSF mitochondrion. The average length of the mature fully-edited mitochondrial rRNAs and mRNAs was considered to be 880 nt and 933 nt, respectively (Kirby et al., 2016). With a cost of 5 ATPs for the synthesis of each rNTP (Table 4), efficient recycling of the ribonucleotides once the RNAs are degraded, 2 ATPs for recharging each monomer (Lynch and Marinov, 2015), and similar half-lives to those RNAs encoded by the nuclear genome, we calculated that 17.3 × 10^7^ and 0.3 × 10^7^ ATP molecules are necessary to synthesize the estimated pool of mitochondrial rRNAs and mRNAs, respectively. In the polycistronic model of transcription, intergenic regions are transcribed and, after RNA processing, the rNTPs are recycled. For that reason, it is necessary to estimate the polymerization cost of the intergenic regions of the polycistrons transcribed from the maxicircles. Given the size of each maxicircle (∼23 kb) and the sum of the average length of mature RNAs (18,554 nt) we considered that 4,446 nt are polymerized for each maxicircle, resulting in a consumption of ∼2.7 × 10^5^ ATP molecules.

To have a more complete estimation of the total transcriptional cost of the mitochondrial genome, it is necessary to estimate the cost of the transcription of gRNAs. Transcription of the minicircles generates an 800 nt precursor (Aphasizhev and Aphasizheva, 2011), encoding 2-5 gRNAs each, with an average length of 49 nt (Cooper et al., 2022). It means that, on average, for each minicircle, 678 rNTPs are polymerized and then recycled after processing. Considering the number of 6,000 minicircles per cell (Cooper et al., 2019) and that at least one of each gRNA will be transcribed, the minimal cost for minicircles transcription is the cost of the polymerization of the rNTPs of the intergenic regions, which is ∼0.8 × 10^7^ ATP molecules, plus the cost of synthesis and polymerization of the rNTPs in the mature gRNAs, which is ∼0.4 × 10^7^ ATP molecules. Thus, transcription of the minicircles costs, at minimum, 1.2 × 10^7^ ATPs per cell cycle.

Assuming that that the transcription of maxicircles has a similar global dynamic as that of nuclear transcription, and that each minicircle is only transcribed once per life cycle, we calculated the cost of transcription of the mitochondrial genome at 18.8 × 10^7^ ATP molecules. It is worth mentioning that these are likely underestimations due to the scarce knowledge of the ATP expenditure of each process involved in kDNA transcription, pre-RNA processing and mRNA editing.

### Energy expenditure for proteome synthesis, maintenance, degradation

Regarding the biosynthesis of proteins, we must take into account the cost of obtaining their components, the amino acids. For this, we consider two sources for these metabolites: their uptake from the environment, and their biosynthesis *de novo*. The present work is based on data obtained by culturing the parasites in a very rich medium containing all the amino acids, so in this condition, and probably also *in vivo* in the bloodstream, it is reasonable to assume that most of their requirements are fulfilled through their acquisition from the extracellular medium. However, we made also an estimation of the cost of the *de novo* synthesis for those amino acids having their biosynthetic pathways predicted from the genome sequence as this estimation could be of general interest (see Supplementary Text 1).

To determine how much ATP is spent by BSF *T. brucei* on protein synthesis, we first estimated the number of amino acids present in its proteome from the cell’s known volume and the calculated protein density. The volume of *T. brucei* BSF (1K1N, *i.e*. one kDNA network and one nucleus, after cell division, before DNA replication) cells is ∼45 µm^3^ (Rotureau et al., 2011). According to the method proposed by Milo (2013), we calculated the number of proteins per cell based on the protein mass per unit volume (*c*_p_) in g of protein per ml of cell volume, which has been estimated for several cell types as being 0.2 g/ml (Albe et al., 1990; Milo, 2013). Other relevant parameters taken into account are the average length of proteins (*l*_aa_) (300 amino acids according to Milo and Phillips, 2015), and the average molecular mass of amino acids (*m*_aa_) (110 Da). Therefore, the average mass of proteins per unit volume (N/V) is:

N/V = *c*_p_ / *l*_aa_ × *m*_aa_ = 6.1 μg/ml (Milo, 2013).

For converting these values into the number of proteins per μm^3^, we applied the following equation:

N/V = (*c*_p_ × N_A_ × 10^-12^ ml/μm^3^) / *l*_aa_ × *m*_aa_

where N_A_ is Avogadrós number. The obtained value is 3.5 × 10^6^ proteins/μm^3^. Therefore, considering a cell volume of 45 μm^3^ we obtained a number of proteins per cell of 157.5 × 10^6^.

With an average protein length of 300 amino acids (Milo and Phillips, 2015), we then calculated that a single cell contains 4.7 × 10^10^ amino acids as protein components (in other words, forming peptide bonds). The direct cost of polymerization is 4 ATPs per amino acid (Mahmoudabadi et al., 2019), so the direct cost of translation, for a single cell, is about ∼1.9 × 10^11^ ATPs to double the entire set of proteins (Table 7).

**Table 6.** Summary of ATP costs per cell cycle associated with kDNA transcription.

| Process | Maxicircles | Minicircles |
| --- | --- | --- |
| Transcription | $17.6 \times 10^7$ | $0.4 \times 10^7$ |
| Polymerization of intergenic regions | $0.03 \times 10^7$ | $0.8 \times 10^7$ |
| **Total** | **$17.6 \times 10^7$** | **$1.2 \times 10^7$** |

**Table 7.** Summary of ATP costs associated with protein synthesis and degradation during a cell cycle of BSF *T. brucei*.

| Process | ATP cost |
| --- | --- |
| Proteome doubling | $1.9 \times 10^{11}$ |
| Protein degradation | $0.18 \times 10^{10}$ |
| Protein resynthesis | $1.4 \times 10^{11}$ |
| **Total** | **$3.48 \times 10^{11}$** |

During the BSF trypanosome’s cell cycle, part of its proteins has to be degraded and replaced by new proteins to be synthesized. The balance between these processes represents the cell’s protein turnover. Its cost must be added to that of the entire proteome doubling during the parasite’s growth and division. We considered for our calculations only regulated protein degradation, which requires an expenditure of 100 – 200 ATP molecules per degraded protein (Lynch and Marinov, 2015; Peth et al., 2013). Here we assumed an average value of 150 ATPs per degraded protein. A proteomic turnover study determined that this process is directly influenced by the duration of the cell cycle. For this, the duration of BSF trypanosomes life cycle was determined as being 11.85 h. This remarkable difference with the duration considered in our study can be explained by the fact that the authors performed this estimation for parasites growing under protein labelling conditions (data were obtained using SILAC labeling for proteomics). Under these conditions, the estimated a half-life for the entire proteome was 5.6 h (Tinti et al., 2019). As we are using, in this work, the duration of the BSF cell cycle of 5.3 h, we made an estimation of energy cost of the proteome’s turnover in our model by scaling the half-life using the rationale described in Tinti et al. The obtained value for the proteome half-life was then 2.56 h, meaning that, according to the exponential decay law, during an entire cell cycle 0.76% of the proteome is degraded. Therefore, 1.2 × 10^8^ proteins per cell are degraded during a cell cycle, at an average cost of 1.8 × 10^10^ ATP molecules (Table 7). At the same time, to maintain the entire proteome, the same quantity of these proteins must be newly synthesized at a cost of 1.4 × 10^11^ (Table 7). This, added to the synthesis of an extra new proteins for obtaining an entire proteome for each daughter cell, requires 3.3 × 10^11^ ATP molecules per cell cycle for protein synthesis. In summary, the total cost for degradation, resynthesizing and doubling of the proteome is ∼3.5 × 10^11^ ATP molecules (Table 7).

### Energy cost of sugar nucleotides used in the synthesis of the VSG coat

In the BSF of *T. brucei*, the major surface protein is the VSG, which is highly glycosylated. The VSG polypeptide is estimated as being present in 10^7^ copies per cell, representing approximately 90% of cell surface polypeptides and 10% of total cellular protein content (Grünfelder et al., 2002). Therefore, the sugar nucleotides used in the synthesis of the VSGs require by far the major part of the ATP dedicated to the synthesis of the entire pool of sugar nucleotides in these cells. Trypanosomatids’ survival, infectiousness, and virulence in their mammalian hosts are directly influenced by their cell surface glycoconjugates. The amount of sugar nucleotide used for their synthesis was calculated based on previous estimates (Turnock and Ferguson, 2007). For this, certain conditions were assumed: i) the metabolites are evenly distributed throughout the cell volume; ii) the demand for each sugar nucleotide is minimal and for this calculation we did take into account the glycoconjugates turnover; iii) the contributions of low-abundance glycoconjugates are considered negligible; and iv) an average glucidic composition of Man_15_GlcNAc/GlcN_5.5_Galp_5_ (Grünfelder et al., 2002), based on that of VSG variant 221 (MITat 1.2). On these bases, we estimated the need for 5 × 10^7^ UDP-Galp, and the same quantity of UDP-GlcNAc. Also, 15 × 10^7^ units of GDP-Man are required. Considering an average ATP expenditure of 4 HEBs (high-energy bonds) per nucleotide sugar, the total ATP requirement for synthesizing the glucidic moieties of 10^7^ VSGs is ∼1 × 10^9^ ATP molecules per cell during a cell cycle (Table 8).

**Table 8.** Estimation of ATP cost for the synthesis of sugar nucleotides for *T. brucei* VSGs per cell cycle.

| Sugar Nucleotide | HEBs per molecule | SN in VSG | HEB per SN | HEB per cell |
| --- | --- | --- | --- | --- |
| UDP-Galp | 4 | 5 | 20 | 2 x 10<sup>8</sup> |
| UDP-GlcNAc | 4 | 5 | 20 | 2 x 10<sup>8</sup> |
| GDP-Man | 4 | 15 | 60 | 6 x 10<sup>8</sup> |
|  |  |  | **Total** | **1 x10<sup>9</sup>** |

### Energy expenditure for doubling the lipidome of *T. brucei* BSF

One of the basic needs for cell proliferation is the production of a new set of lipids for synthesizing the external and internal membranes. However, we have not found in the literature an estimate of the total energy cost necessary for doubling the total cell membrane content. BSF *T. brucei* can obtain its lipids by two different routes (Paul et al., 2001; Poudyal and Paul, 2022): either from the mammalian host plasma, mainly by receptor-mediated endocytosis of LDL particles (Coppens et al., 1995) or by *de novo* synthesis. The contribution of both routes may vary dependent on external conditions. A calculation of the cost of lipid acquisition by uptake from the host is an integral part of the estimation of the total cost of the formation of endocytic vesicles described below. Given that *T. brucei*’s total pool of phospholipids and sterols (Guan and Mäser, 2017; Patnaik et al., 1993; Richmond et al., 2010), as well as their biosynthesis pathways (Dawoody Nejad et al., 2020; Gibellini et al., 2008; Lee et al., 2006; Lilley et al., 2014) have been characterized in detail, it allowed us to estimate the energy requirements when doubling of the lipidome content of BSF trypanosomes would entirely occur by *de novo* routes. For this purpose, we considered the number of HEBs used in the biosynthetic pathways of each species of phospholipid and ergosterol. With this information, we were able to estimate the amount of ATP needed for their doubling (Table 9).

**Table 9.** . Lipid composition and energy cost of biosynthesis for each molecular species in BSF of *T. brucei*.

| Species | %mol | [Conc]<br>(nmol.<br>mg<sup>-1</sup>) | Number of<br>HEB for<br>biosynthesis | nmol<sub>HEB</sub><br>ug prot | nmol<sub>HEB</sub><br>per<br>parasite | ATP<br>molecules<br>per<br>parasite |
| --- | --- | --- | --- | --- | --- | --- |
| PC | 47.8 | 171.6 | 4 | 686.4 | 6.86 x 10<sup>-6</sup> | 4.1 x 10<sup>9</sup> |
| PE | 20.7 | 74.313 | 4 | 297.252 | 2.97 x 10<sup>-6</sup> | 1.8 x 10<sup>9</sup> |
| PI | 5.4 | 19.386 | 4 | 77.544 | 7.75 x 10<sup>-7</sup> | 4.7 x 10<sup>8</sup> |
| PS | 3 | 10.77 | 4 | 43.08 | 4.31 x 10<sup>-7</sup> | 2.6 x 10<sup>8</sup> |
| CL<sup>1</sup> | 0.715 | 2.56 | 8 | 20.48 | 2.05 x 10<sup>-7</sup> | 1.2 x 10<sup>8</sup> |
| PG<sup>1</sup> | 0.485 | 1.74 | 4 | 6.96 | 6.96 x 10<sup>-8</sup> | 4.2 x 10<sup>7</sup> |
| Ergosterol<sub>2</sub> | 13.8 | 49.54 | 12 | 594.48 | 5.94 x 10<sup>-6</sup> | 3.6 x 10<sup>9</sup> |
| SM | 14.5 | 52.055 | 4 (5 for IPC<br>or EPC) | 247.7 | 2.48 x 10<sup>-6</sup> | 1.5 x 10<sup>9</sup> |
| **Total** |  |  |  |  | **1.97 x 10<sup>-5</sup>** | **1.19x10<sup>10</sup>** |

PC, phosphatidylcholine; PE, phosphatidylethanolamine; PI, phosphatidylinositol; PS, phosphatidylserine; CL, cardiolipin; PG, phosphatidylglycerol SM, sphingomyelin. ^1^ Based on the molar fraction of PG/CL found in the procyclic form; ^2^ compositions were observed in neutral fractions. E.U., Elementary Units.

The total amount of ATP consumed during the life cycle of *T. brucei*, for the entire lipidome doubling (which includes the cost of membrane doubling) is 1.19 × 10^10^ ATP molecules. Noteworthy, among the costs calculated (Table 9), the species that are most energy demanding are ergosterol (36%), PC (31.6%), PE (13.7%) and SM (12%), respectively.

### Energy expenditure on polyphosphate synthesis

Polyphosphates (polyP) are linear polymers of a few to many hundreds of inorganic phosphate (Pi) residues linked by HEBs. They are arbitrarily divided into two forms: short-chain (SC, from 3 to ∼300 Pi) and long-chain (LC, from 300 to ∼1000 Pi), based on the method used for their extraction (Moreno and Docampo, 2013). In trypanosomatids, the polyP has been proposed to be associated with several biological functions, such as osmoregulation (Docampo et al., 2010), Ca^2+^ signaling (Lander et al., 2016) and energy source storage (Docampo et al., 2010). Most polyPs in trypanosomatids are concentrated in acidocalcisomes (Docampo et al., 2010), although polyP has also been found in the nucleus, cytosol and glycosomes. However, in BSF, polyPs have been detected mostly in acidocalcisomes and glycosomes (Negreiros et al., 2018). PolyP is very abundant in BSF: 600 μM for LC and 250 μM for SC for 2 x10^7^ parasites (Lemercier et al., 2004). As the amount of polyP is measured by the molarity of phosphate units, these concentrations correspond to the number of monomers in the polymerized inorganic phosphates (Cordeiro et al., 2019). So, we consider LC+SC as the total concentration of polyP corresponding to 850 μM for the extraction from 2 × 10^7^ parasites. Based on a cellular volume of 45 μm^3^ per individual cell, we obtained a total volume of 0.9 μL for 2 x10^7^ BSF cells. The 850 μM of Pi polymerized in polyP in 0.9 μL corresponds to 765 × 10^-12^ mol for 2 × 10^7^ parasites. As each Pi corresponds to one HEB, which is equivalent to one ATP molecule, the total ATP required to synthesize the BSF’s whole content of polyP is 3.8 × 10^-17^ mol of ATP, in other words, 2.3 × 10^7^ ATP molecules per parasite. Knocking out the Vacuolar Transporter Chaperone 4 in *T. brucei* caused a decrease of 25% of the total polyP (Lander et al., 2013). As BSF are not challenged by strong osmolarity or nutritional variations (the main processes in which polyP are spent) (Lander et al., 2013), we assume that this is the global rate of polyP degradation. During a cell cycle, BSF has to synthesize at least a new set of polyP for replication and renew the 25% of the polyP stock. Thus, the total ATP demand for synthesizing a new set of polyPs and maintaining the existing one is 2.9 × 10^7^ molecules. Additionally, Lander et al. (2013) showed that each Pi translocated into the acidocalcisomes requires the consumption of one ATP molecule (Lander et al., 2013). However, the transport of small molecules such as Pi into the glycosomes is not expected to have any cost since low molecular mass molecules and ions can freely diffuse through pores in the glycosomal membrane (Gualdron-López et al., 2012). Considering these facts, and that almost all polyP is stored in acidocalcisomes (Docampo et al., 2010; Negreiros et al., 2018), we can calculate the number of ATP molecules to transport into these organelles the Pi units needed for polyP polymerization. This implies the same number of Pi units imported into the acidocalcisomes as those used for polymerization, which doubles the budget resulting in total consumption of 5.8 × 10^7^ ATP molecules per cell cycle.

### Vitamins and other micronutrients

Trypanosomes also need vitamins and other micronutrients whose biosynthetic processes and/or uptake require ATP. Mechanisms for uptake from the medium have been identified for choline (Macêdo et al., 2013), pyridoxine (vitamin B6) (Gray, 1995) and riboflavin (vitamin B2) (Balcazar et al., 2017). Ascorbic acid (vitamin C) biosynthesis has been identified in *T. brucei*, with the last step taking place within glycosomes (Wilkinson et al., 2005). Vitamin B1 is especially interesting because it is not efficiently taken up under physiological conditions suggesting that its intracellular levels must be obtained via biosynthesis (Stoffel et al., 2006). Overall, considering the nutrients mentioned above, there is still much to be elucidated. Although there is evidence that biosynthesis occurs for some vitamins, such as B1 and B6, the pathways themselves are not understood in detail (Gray, 1995; Stoffel et al., 2006). All works referenced in this section that identified an uptake mechanism for nutrients describe passive processes. Even if some of these compounds are biosynthesized, most of them are produced in low quantities. In summary, there is no evidence that these processes impact ATP levels meaningfully. Our conclusion for now, with some reservations, is that vitamin transport and biosynthesis do not have a significant impact on the energy budget of the parasite.

### ATP requirement for transmembrane transport

The cellular uptake of molecules and ions is part of the cell maintenance processes, and in most cases, it has an energy cost (Lynch and Marinov, 2017). The energy dedicated to cell maintenance includes a contribution necessary for preserving a homeostatic ionic composition (Stouthamer and Bettenhaussen, 1973). The energy demand by the uptake of amino acids, ammonium, potassium ions and inorganic phosphate from the extracellular medium into the cell was previously estimated for the synthesis of a new microbial cell, *in casu Escherichia coli* (Stouthamer, 1973). To obtain a value for the energy demand of BSF transport processes, we used the calculations made by Stouthamer as a model (Table 10). Stouthamer assumed that 0.5 moles of ATP are necessary for the uptake of 1 mole of NH4^+^, and 1 mole of ATP is necessary for the uptake of 1 mole of Pi, any amino acid, acetate or malate. For Na^+^ and K^+^ data are available for BSF *T. brucei*, allowing us to make a quite accurate calculation, and the cost of moving them across the plasma membrane was estimated separately (see below). It is worth mentioning that Stouthamer did not consider the costs of taking up glucose, which could be relevant for many prokaryotes but not for *T. brucei* where glucose transport happens by facilitated diffusion (J Gruenberg et al., 1978; ter Kuile and Opperdoes, 1991b). For *E. coli*, depending on the culture conditions, between 18.3 and 19.4% of the total energy required for a cell formation is needed only for overall solutes uptake (Stouthamer, 1973). Due to the lack of other data, we considered that BSF of *T. brucei* uses an intermediate percentage of its total ATP budget for solutes uptake (18.9%), representing ∼1.1 × 10^11^ ATP/ cell cycle × cell. For the calculation of costs of the transport of Na^+^ and K^+^, we used data previously obtained (Bridges et al., 2008; Nolan and Voorheis, 2000). Considering that the ouabain-sensitive BSF Na^+^/K^+^ ATPase has a specific activity for ATP hydrolysis of ∼1.17 nmoles/min × mg (equivalent to ∼1.17 nmoles/min × 10^8^ cells (Opperdoes et al., 1984)) and that ATP is hydrolyzed into ADP + Pi with the concomitant exchange of 3 Na^+^ for 2 K^+^, we calculated that a continuous activity of this pump during 5.3 hours would result in an ATP cost of 2.2 × 10^5^ ATP molecules per cell during an entire cell cycle. This value is negligible when compared to the total cost of transport of other ions and metabolites. Additionally, H^+^-ATPase is important to regulate the intracellular pH of BSF *T. brucei* and an approximate value of 534 nmol/min × mg protein was reported for the H^+^ efflux (Vanderheyden et al., 2000). Taking account of this value and the Stouthamer assumptions, we estimated that ∼1.02 × 10^10^ ATP molecules are necessary for the H^+^ efflux per cell during an entire cell cycle. Regarding Ca^2+^ efflux, proteins with homology to PMCA-type Ca^2+^-ATPases were identified and reported in *T. brucei* as TbPMC1 and TbPMC2 (Luo et al., 2004). In particular, TbPMC1 has been located in the plasma membrane of BSF. However, no information is available regarding its ATP consumption. Even so, we suggest that compared with the values estimated by Luo et al., the ATP expenditure for Ca^2+^ efflux could be negligible when compared to the total cost of transport in the parasite. Based on these calculations, the estimated ATP costs for transport of solutes across the plasma membrane were estimated as being ∼1.2 × 10^11^ ATP/ cell cycle × cell.

**Table 10.** ATP requirement for the formation of microbial cells from glucose and inorganic salts and the influence of the addition of amino acids (AA) or/and nucleic acid bases (bases). (Modified from Stouthamer, 1973).

| Ion/Metabolite | ATP<br>required<br>(moles x $10^{-4}$<br>/ g cells) | % of the total | ATP required<br>(moles x $10^{-4}$ /<br>g cells) | % of the total |
| --- | --- | --- | --- | --- |
|  | AA | AA | AA + bases | AA + bases |
| $NH_4^+$ | 10.4 | 3.0 | 0.0 | 0.0 |
| Amino acids | 47.9 | 13.7 | 47.9 | 15.3 |
| Phosphate ions | 7.7 | 2.2 | 7.7 | 2.5 |
| **Total** | **66** | **18.9** | **55.6** | **17.7** |

### The cost of motility

Motility due to beating of its single flagellum serves the trypanosome to navigate the environment. But for BSF *T. brucei* it has the important additional role of counteracting the defense of the infected host, as it enables clearance of host antibodies attached to VSGs by causing these surface coat proteins to be recycled (Engstler et al., 2007). As a curiosity, the name *Trypanosoma* is derived from the Greek word describing the peculiar movement of these cells (auger cells) (Shimogawa et al., 2018). Trypanosomes are vigorous swimmers, and the swimming velocity depends on the microenvironment’s viscosity. They can reach a speed of at least 20 μm/s, allowing the hydrodynamic removal of attached host antibodies (Heddergott et al., 2012). The frequency of flagellar beating has been measured as 15-20 Hz (Stellamanns et al., 2014). Considering that the resultant energy during the breakdown of 1 ATP molecule is 5.064 × 10^-20^ J and that the power generated by a flagellar beating is ∼4 × 10^-17^ J, one flagellar beating results from the consumption of at least 790 ATP molecules. If we assume that the ATP hydrolysis for flagellar motility is constant, based on the speed maintenance (output) and on the fact that trypanosomes are non-stopping engines, the total ATP consumed can be calculated as:

ATP_f_ = F × n × t

where ATP_f_ is the amount of ATP consumed by the flagellar movement during the entire BSF cell cycle, F is the frequency of flagellar beating (the average value of 17.5 Hz was taken for this calculation), n is the number of ATP molecules consumed per flagellar beating and t is the duration of the cell cycle in seconds. This calculation points out that permanent flagellar beating consumes 2.6 × 10^8^ ATP molecules per cell per cell cycle. This calculation does not take into account the specific characteristics of the internal flagellar machinery, which is responsible for transducing the energy obtained from ATP breakdown into flagellar beating. Inside a flagellum, the axoneme is constituted by 96 nm dynein repeats, forming two central double microtubules surrounded by nine other pairs of microtubules (9(2) + 2) (Ralston and Hill, 2008). The basic dynein composition of each repeat is five outer arms (two-headed) and seven single-headed inner arms of dyneins (Imhof et al., 2019). Each dynein head has an AAA-ATPase domain (Trott et al., 2018), so in total, the axoneme has 17 ATPase domains at each 96 nm dynein repeat. As there are 2 × 9 microtubules in a flagellum, there are a total of 306 ATPase domains/repeat. The average length of a BSF flagellum is 25.3 μm (Imhof et al., 2019), so dividing it by 96 nm, it is possible to calculate that a *T. brucei* flagellum has approximately 264 repeats with ∼40,392 dynein molecules. However, not all components of the flagellar machinery work at the same time. In order to generate a planar waveform, only some of the doublets are activated simultaneously, and the activity should switch periodically between two nearly-opposed doublets (Chen et al., 2015). Considering that each beat is nearly planar in *T. brucei* BSF, instead of more than 40,392 dyneins operating at the same time, there will be those corresponding to 2 out of 9 pairs working simultaneously, in other words, 8,976 active dynein molecules per beat. To estimate the ATP consumption based on the dynein number, it must be considered that every single conformational change in dynein is driven by the formation of an ATP-dynein complex, which is before the power stroke. The power stroke is the motor force that drives the sliding displacement on the longitudinal axis of an axoneme (Lin et al., 2014). The product of axonemal diameter and the shear angle (defined as the interior angle between the symmetry axis of the dynein head and the line tangent to the axoneme, immediately after the first bend), gives the total sliding displacement along an axoneme between two neighboring doublets (Brokaw, 1989; Chen et al., 2015). For *Chlamydomonas*, it was established that the shear angle is ∼1 rad. The diameter of an axoneme is ∼150 nm (Bastin et al., 2000; Höög et al., 2014; Koyfman et al., 2011). The dynein sliding displacement has been calculated as being 8 nm (Lin et al., 2014). As a result, we have the ratio between the sliding displacement and the dynein power stroke, which results in 19 steps per flagellar beat. Assuming that each dynein takes 1 ATP per step, 8,976 of the dynein molecules being active at a given time, and that a flagellar beat needs 19 dynein steps along the microtubules, the parasite has to invest ∼1.7 × 10^5^ ATP molecules per flagellar beat. As previously stated, the average frequency for flagellar beating is 17.5 Hz. Remaking the calculation above with data from the mechanistic analysis of the flagellar machinery (see equation above) the ATP demand by the whole flagellar machinery would be equivalent to 5.7 × 10^10^ molecules.

### ATP cost of activation and recruitment of vesicles

Endocytosis is a very important biological process in *T. brucei*, to capture specific compounds from the environment, such as low-density lipoprotein containing lipids and transferrin providing iron by the receptor-mediated process and serum proteins like albumin complexed with various molecules by fluid-phase endocytosis (Coppens et al., 1995, 1988; Kariuki et al., 2019). However, the mechanisms involved in this process have still not been fully described in this parasite (Link et al., 2021). The process is also crucial for the above-mentioned antibody clearance and VSG recycling which allows the trypanosome to escape from the host immune attack (Manna et al., 2014). BSF possesses at least 10^7^ VSG molecules per cell and recycles the entire VSG coat each 12 min (Engstler et al., 2007). To recycle VSGs, *T. brucei* depends on both endocytic and exocytic pathways. The VSGs are returned to the surface after passing through endosomes where any attached antibodies are removed and routed to the lysosomes for degradation. As every endocytic event in *T. brucei,* it depends on clathrin. For that, the cell produces 6-7 clathrin-coated vesicles bearing VSGs per second (Engstler et al., 2004). For our calculation, we used the minimum value of 6 clathrin-coated vesicles bearing VSGs per second, which implies that these cells would be internalizing 21,600 vesicles per hour. Vesicle formation for VSG recycling is a Rab11-dependent process (Grünfelder et al., 2003). Considering that the cell produces 21,600 vesicles per hour and at least 1 Rab assembly is necessary for each vesicle, the energy cost for the activation and recruitment of vesicles based on the assembly of Rab proteins (that use 1 GTP/Rab) results in a cost of 1.14 × 10^5^ GTP molecules per cell cycle. In this calculation, we are not taking into account some other processes that could impact endocytosis-related ATP consumption in *T. brucei*. Even though the endocytosis process is quite well understood in other organisms such as several opisthokonts (Adung’a et al., 2013), we are not able to estimate other ATP expenditures that can contribute to the total cost of endocytosis in *T. brucei* due to the low conservation of components for this machinery. For example, during the formation of the vesicles, several proteins are recruited to the site of membrane bending, and an actin bridge is built up (Paraan et al., 2020). The assembly of these proteins and the actin polymerization surrounding the vesicle involves ATP and GTP hydrolysis and/or cycling. For cargo translocation along tubulin microtubules, cycling of GTP is also necessary (Kaksonen and Roux, 2018). Clathrin and adaptor protein release also depends on ATPase activity (Hannan et al., 1998). Therefore, the minimal amount of ATP consumed for this process, considering a rate of conversion of 1 ATP per GTP is 1.14 × 10^5^ ATP molecules per cell cycle.

### How much ATP hydrolysis is required to maintain the mitochondrial inner membrane potential (ΔΨm)?

The single mitochondrion of BSF *T. brucei* displays marked differences when compared to those of every other eukaryote described so far, and even when compared to that of other life cycle stages of the parasite, such as the procyclic form. The most remarkable differences are: i) the absence of OxPhos; ii) a marked reduction in the expression levels of proton pumping respiratory enzyme complexes; and iii) a drastic reduction in the expression of TCA cycle enzymes (Zíková et al., 2017). As the mitochondrial integrity and biogenesis depends on the mitochondrial inner membrane potential (ΔΨ_m_) (Brown et al., 2006; Schnaufer et al., 2005), BSF compensates for the lack of functional respiratory proton pumps by using the F_1_F_o_-ATP-synthase in reverse mode. In this way, ΔΨ_m_ is built up and maintained by pumping protons into the intermembrane space by hydrolysis of ATP (Brown et al., 2006). Additionally, the cells require intramitochondrial ATP to prevent inhibition of the trypanosome alternative oxidase, which is needed to use oxygen as a terminal electron acceptor (Luévano-Martínez et al., 2020). It must be noted that the ATP required in the mitochondrial matrix to keep both systems working does not necessarily depend on ATP import by the adenine nucleotide carrier. In the absence of this transporter’s activity, it can also rely on an intramitochondrial substrate-level phosphorylation system, comprising the acetate:succinate CoA transferase and the succinyl-CoA synthetase (ASCT/SCS) cycle. This is reminiscent to the substrate-level phosphorylation and reversal of the ATP-synthase shown in other systems such as the isolated liver and heart rabbit mitochondria (Chinopoulos et al., 2010). Such a system has been demonstrated as being functional in BSF in terms of intramitochondrial ATP production (Jenkins et al., 1988). We hypothesize that this mitochondrial substrate-level phosphorylation system is the main source of intra-mitochondrial ATP, and it can provide sufficient ATP to maintain the ΔΨ_m_ (Millerioux et al., 2012; Mochizuki et al., 2020), despite its relatively low capacity for producing only small quantities of ATP (Michels et al., 2021). In terms of energy expenditure, the mitochondrial substrate-level phosphorylation could then be considered energetically neutral since all ATP produced by the ASCT/SCS cycle is devoted to the maintenance of the mitochondrial membrane potential generated by the F_1_F_o_-ATP synthase.

## Discussion

Long-slender bloodstream forms of *T. brucei* have a unique configuration in terms of the bioenergetic pathways responsible for ATP production. Despite having a mitochondrion, these cells rely almost exclusively on glycolysis for ATP production, and they are the only case in nature (to the best of our knowledge) of mitochondriated cells having a mitochondrion that, under the conditions studied thus far, does not contribute to the cell’s net ATP production. Even more, their ATP synthase hydrolyzes ATP to maintain the mitochondrial inner membrane potential (Nolan and Voorheis, 1992; Panicucci et al., 2017; Schnaufer et al., 2005). Based on data available in the literature on glycolytic flux during proliferation (Haanstra et al., 2012), we calculated with some precision the total amount of ATP produced during a BSF cell cycle, in other words, we calculated the ATP necessary for maintaining alive a BSF trypanosome and building a new one (∼6 × 10^11^ molecules) when cultured in the rich medium HMI-9 medium. There is not much information available on the ATP necessary for the survival and replication of other cell types. In fact, to the best of our knowledge, such numbers have been reported so far only for two other eukaryotic cells and two prokaryotic cells. When our numbers are compared with those of the other cells (Table 11) it becomes clear that, as expected, the ATP produced per cell during a cell cycle is much lower than that for mammalian cells. However, it should be noted that, equally expected, this value is much higher than that obtained for prokaryotic cells.

**Table 11.** ATP produced during the cell cycle in different cells.

| Organism | Total | Reference |
| --- | --- | --- |
| *Mycoplasma pneumoniae* | $4.3 \times 10^9$ | (Wodke et al., 2013) |
| *Escherichia coli* | $5.9\text{-}12 \times 10^9$ | (Farmer and Jones, 1976;<br>Stouthamer, 1973) |
| BSF *Trypanosoma brucei* | $5.94 \times 10^{11}$ | This work |
| Mammalian tissue culture cell iBMK | $1.2 \times 10^{13}$ | (Fan et al., 2013) |
| Human fibroblast | $4.5 \times 10^{13}$ | (Flamholz et al., 2014) |

Once knowing how much ATP is available for keeping alive and replicate a cell, it was interesting to analyze how much of this valuable resource is used for critical biological processes (Figure 1, Supplementary Table S5). The cost of DNA replication depends, in addition to the genome size, on the nucleotide composition and the specific ATP cost of their biosynthesis. Whilst in other organisms the average cost spent on dNTP biosynthesis from glucose is 50 ATP molecules (Lynch and Marinov, 2015), in *T. brucei* BSF it is only 10 ATP molecules. This is due to the fact that this parasite does not synthesize purines *de novo* but uses the salvage pathway, and synthesizes pyrimidines from externally supplied glutamine and aspartate. According to our calculations, 90% of the costs of the total DNA duplication is the cost of replicating the nuclear genome, while the remaining 10% corresponds to the cost of replicating the kDNA. Differently from replication, transcription costs are strongly influenced by other factors. Large polycistronic units are often assumed as costly because they involve the transcription of “useless” DNA (for example intergenic sequences, developmentally regulated genes, pseudogenes, *etc*.) that must be further eliminated during the trans-splicing processing for producing the mature mRNA, or post-transcriptional degradation. However, according to our calculations, a significant part of the cost of transcription is due to the biosynthesis of rNTPs. As rNTP used in transcribing the intergenic regions can be recycled they would not constitute an extra cost (Lynch and Marinov, 2015). So, the only extra cost that can be assumed is that of their polymerization (equivalent to 2 ATP/base). Considering this, the extra cost of polycistronic transcription is ∼30% of the total transcriptional cost, in other words, 0.3% of the total budget for maintaining and building a new cell. There are not enough data to calculate in detail the total extra cost of transcribing coding sequences that must be further degraded in order to control gene expression. However, some estimations can be made based on the fact that only 47 out of 9,694 (∼0.5%) genes are considered as not being expressed in BSF, and 772 out of 9,694 (8%) genes are considered down-regulated in BSF when compared to procyclic forms (Naguleswaran et al., 2018). Considering the extreme case in which both gene populations are completely degraded after polymerization, the spurious coding RNA polymerization corresponds to 8.5% of the total coding RNAs. As we considered 2 ATP molecules being spent per base polymerized, an average transcript length of 2,800 nt, and an average RNA synthesis of 1.2 RNAs/h (estimated in Haanstra et al., 2008) the estimated ATP expenditure is ∼2.9 × 10^7^ ATP molecules (0.5% of the total ATP expenditure for the completion of a cell cycle) (Figure 1, Supplementary Table S5). These values can be compared with those that can be estimated from a scenario of having transcriptional regulation for each protein-encoding gene. Considering that BSF expresses 8,875 genes this implies the formation of at least an equivalent number of transcriptional OCs, instead of the reduced number of OCs necessary in the polycistronic transcription system. Based on an individual cost of 20 ATP molecules/OC, the total cost of individual transcriptional initiation would be 1.7 × 10^5^ ATPs, a much higher value when compared to the ∼1 × 10^4^ ATPs required for OCs in the polycistronic transcription. Regardless of the case, both costs seem to be largely negligible concerning the total transcription cost and therefore from a purely energetic point of view the evolutionary advantage of individual transcription seems to be impactless.

**Figure 1.**
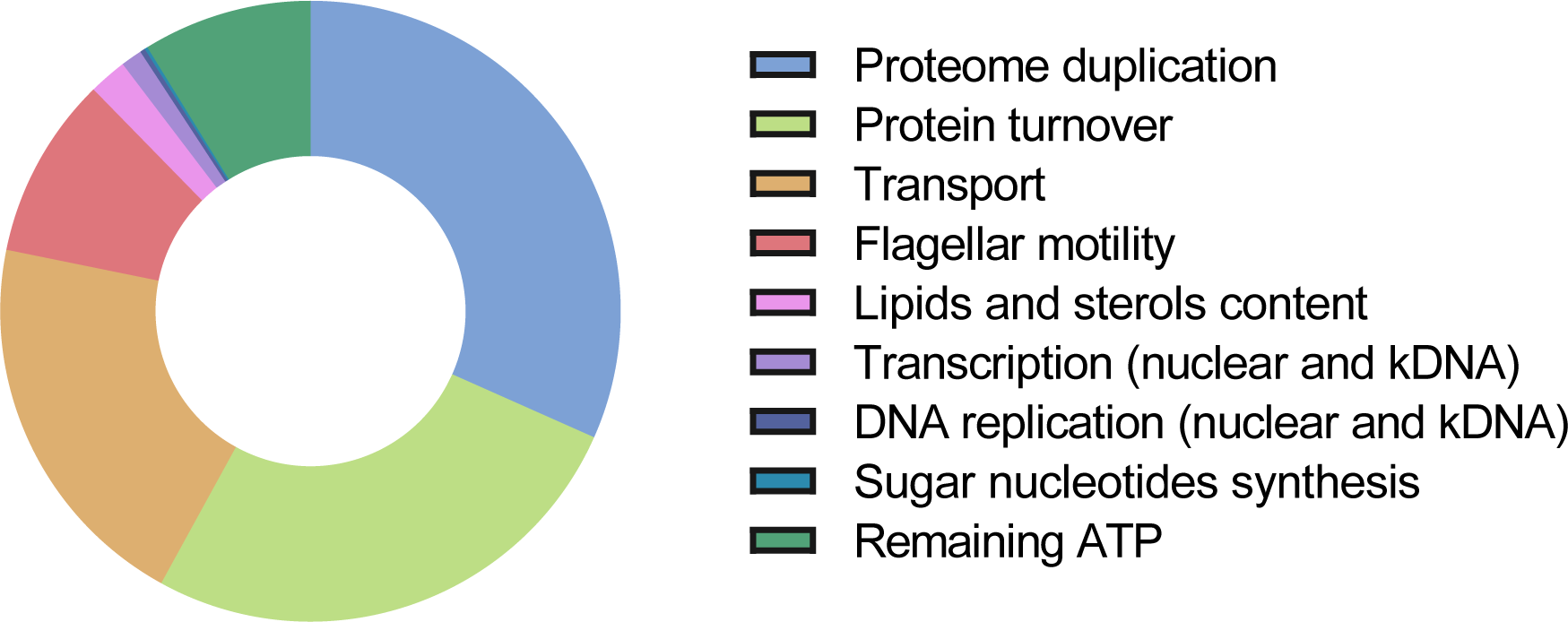
Summary of the most energetically costly biological processes in bloodstream form *T. brucei.* For underlying values see text and Supplementary Table S5.

As reported for several cell types, the synthesis and maintenance of the proteome is the most expensive process during a cell cycle (Supplementary Table 5 and Table 12). Despite the fact that BSF trypanosomes take up most of the amino acids from the medium instead of *de novo* synthesizing them, they are, according to our calculations, not an exception with regard to the expensiveness of proteome production and maintenance. This is explainable because the formation of peptide bonds is one of the costliest biochemical reactions in a cell (4 ATP molecules per bond). Therefore, taken together, translation and protein turnover demand 58.6% of the ATP budget (Figure 1). An interesting point emerges when analysing the cost of synthesizing the amino acids that compose the proteome in comparison with the energy required to import them from the environment. According to Mahmoudabadi, the average cost of synthesizing 1 amino acid is 2 ATPs (Mahmoudabadi et al., 2019). We are assuming that during proteome turnover all amino acids are recycled. Thus, cost of synthesizing will only be considered for amino acids to be used for building a new proteome (not for the maintenance due to turnover). We estimated that the synthesis of a new proteome demands 4.7 × 10^10^ amino acids. Therefore, the cost of synthesizing all amino acids would be 9.4 × 10^10^ ATPs. Herein we assumed that the total cost of uptake of amino acids and ions was as estimated by Stouthamer for *E. coli* (between 13.7 and 15.3% of the total cell ATP budget). Taking the intermediate value of 14.5%, this would result in an ATP cost of 8.6 × 10^10^, surprisingly very close to the cost estimated for amino acid biosynthesis. It is generally assumed that taking metabolites up is energetically more efficient than synthesizing them, and this efficiency would contribute to the parasitic lifestyle. Our calculations show that, in principle, for amino acids, the difference is very minor, impacting the total budget by less than 1.5%. These calculations do not include the cost of the synthesis of sugar nucleotides used for the glycosylation of surface proteins (mostly VSGs). Even being part of the total cost of building an entirely new proteome, it represents a negligible 0.5% of the total ATP demanded by this process.

**Table 12.** Comparison of ATP demand in different cell types.

| Process | ATP demand (%) |  |  |
| --- | --- | --- | --- |
|  | BSF *T. brucei*<br>(This work) | Bacteria (Russell<br>and Cook, 1995;<br>Stouthamer, 1973)<sup>a</sup> | Mammalian cells<br>(Buttgereit and<br>Brand, 1995) |
| DNA replication | 0.3 | 1.8 | 25<sup>c</sup> |
| Transcription | 1.1 | 11.8<sup>b</sup> |  |
| Proteome doubling | 31.7 | 59.3<sup>d</sup> | 34<sup>d</sup> |
| Protein turnover | 26.3 |  |  |
| Sugar nucleotides synthesis | 0.2 | ND | ND |
| Lipids and sterols synthesis | 2 | 0.3 | ND |
| polyP synthesis and maintenance | 0 | ND | ND |
| Transport (aa, K<sup>+</sup>, Pi<sup>-</sup>) | 20.2 | 18.1 | 33<sup>e</sup> |
| Flagellar motility | 9.5 | ND | ND |
| Activation/recruitment of vesicles | 0 | ND | ND |

<sup>a</sup> Bacteria grown in the presence of glucose, inorganic salts and amino acids
<sup>b</sup> Sum of RNA synthesis and turnover
<sup>c</sup> Sum of DNA/RNA synthesis
<sup>d</sup> Reference refers only to protein synthesis
<sup>e</sup> Sum of Na<sup>+</sup>/K<sup>+</sup> and Ca<sup>2+</sup> ATPases
ND, not determined

Regarding the cost of synthesizing the lipidome, it is interesting to note that BSF trypanosomes contain most of the lipids commonly present in eukaryotic cells (Carroll and McCrorie, 1986). Although BSF *T. brucei* can acquire most of the lipids from the blood of the mammalian host (Coppens et al., 1988), they have also the ability to rely on complete *de novo* biosynthesis of phospholipids and glycolipids to fulfill the need of some specific lipids (van Hellemond and Tielens, 2006). For example, the VSG synthesis and anchoring in the plasma membrane requires high quantities of myristate, which is at low abundance in the host serum (Buxbaum et al., 1996). As most of the lipids biosynthesis pathways have been characterized in detail for *T. brucei* (Dawoody Nejad et al., 2020; Gibellini et al., 2008; Lee et al., 2006; Lilley et al., 2014), we could estimate that the synthesis of the complete repertoire of lipids and sterols would consume 2% of the total ATP budget (Table 12, Figure 1). However, this value is likely to be an overestimation, since data indicate a balance between transport and biosynthesis is responsible for the maintenance of the lipids content in BSF *T. brucei* (Poudyal and Paul, 2022).

PolyPs are ubiquitously distributed among bacteria, protists and mammalian cells, and in unicellular eukaryotes have been proposed to have a role in different biological processes such as adaptation to stress, osmoregulation and metabolism regulation. In prokaryotes, they have been proposed as storage of HEBs. Indeed, their hydrolysis involves the possibility of being coupled to phosphorylating ADP to ATP. However, based on our calculations, a role for polyPs as an energy reservoir seems unlikely. In BSF trypanosomes, polyPs are synthesized inside acidocalcisomes, which necessitates the import of Pi units into this organelle, an ATP-demanding process. According to our calculations, this implies the expenditure of 2 ATP molecules per unit of Pi polymerized, in other words, the polymerization requires at least twice the energy that can be retrieved by hydrolysis. This, together with the fact that the total energy stored in the form of polyPs is less than 0.005% of the total ATP produced during a cell cycle (Supplementary Table S6) suggest that their use as an energy reserve could only be restricted to very specific processes.

Regarding the costs of critical processes for survival and replication of BSF not related to the maintenance and duplication of biomass, we estimated the costs of motility, endo/exocytic vesicles formation, and the maintenance of the mitochondrial inner-membrane potential (which in the case of BSF is exclusively dependent on ATP hydrolysis). Motility occurs as a non-stop process during the entire cell cycle and is associated with the activity of the flagellar machinery. Two calculations were made on the basis of data available in the literature: i. based on the energy dissipated by the flagellar beating; and ii. based on the ATP demand of the flagellar structure, relying on the information on the composition and organization of the molecular motors responsible for the flagellar movement. Both calculations resulted in values differing in two orders of magnitude. It must be noted that both values refer to different phenomena since in the first case we estimated the energy output and in the second case the energy demand of the entire flagellar machinery. Therefore, if both values are correct, the efficiency of the machinery for flagellar beating can be calculated as the percentual ratio between the energy output and input, in this case approximately 0.5%. In this sense, it should be pointed out that Stellamanns et al. (2014) found a discrepancy between the power necessary to move the BSF body in relation to that actually produced by the flagellar movement in the range of one order of magnitude(Stellamanns et al., 2014). Whatever the case, the low efficiency of this process in terms of trypanosome motility is in agreement with the fact that flagellar beating is necessary for other processes not necessarily related to parasite movement, such as VSG recycling for antibody clearance (Engstler et al., 2007; Stellamanns et al., 2014). To estimate the total percentage of the budget used for flagellar beating, we considered the highest value obtained, which resulted in the consumption of 9.6% of ATP produced (Figure 1). Regarding the VSG recycling and antibody clearance, they require, in addition to flagellar movement, the formation of vesicles for trafficking the surface proteins through the cell interior. Due to the fact that the ATP (or in some cases GTP) requirements of these processes are largely unknown, we did not consider the cost of formation of the actin bridge, the cargo translocation along tubulin microtubules, and the clathrin and adaptor protein release (Hannan et al., 1998). Therefore, the ATP cost in our calculation is probably underestimated. However, as it represents less than approximately 0.0001%, the whole process is likely to be energetically undemanding.

In this paper, we reported our calculation of the energy budget of maintaining alive and building up a BSF cell of *T. brucei* during its cell cycle based on the cellular and metabolic processes known to occur in these trypanosomes and data available about the ATP costs of the processes. Where relevant data for *T. brucei* were lacking, we estimated the costs based on data known for other organisms. Of course, the outcome of this endeavour is an approximation; for several processes in the trypanosomes, or even in general in cells, quantitative information is not available and/or how much ATP is required to sustain them is unknown. Nonetheless, the approximation seems realistic; all known major processes have been considered. Our analysis provided results that are amenable for experimental interrogation, while it also revealed where more research is required to allow an even more complete understanding of the energy expenditure of trypanosomes. Moreover, it will be interesting to expand this study to the analysis of other proliferative life-cycle stages of *T. brucei*, or those of related parasitic (*e.g.* the intracellular *T. cruzi* amastigote) and free-living organisms for which sufficient data are or may become available in the foreseeable future.

## Materials and Methods

## Databases

## Methods

(1) Our analysis is restricted to long-slender proliferating forms of BSF *T. brucei*. For their energy supply, these trypanosomes are entirely dependent on glucose uptake from the blood. Almost all glucose is converted to pyruvate, which is excreted, resulting in a yield of 2 ATP/glucose consumed. We have based our calculations on the quantitative analysis of the glucose consumption rate in exponentially growing trypanosomes of *T. brucei* strain Lister 427 with a doubling time of 5.3 h, as described by Haanstra et al., 2012. All calculations for rates of ATP consumption in different processes and activities of trypanosomes as described in the literature have been scaled to a cell cycle of 5.3 h.

(2) ATP consumption for biosynthetic processes has been calculated taking into account the (macro)molecular content (proteins, nucleic acids, lipids) of the trypanosomes, the known precursors which are either synthesized or taken up from the host environment, the rate of the processes and the turnover of the (macro)molecules. Also, the energy of uptake of processes was considered.

(3) Other energy costs that were estimated involved: biogenesis of subcellular structures, endocytosis and recycling of the VSG surface coat, motility, protein degradation, and generation and maintenance of transmembrane electrochemical ion gradients.

### Detailed costs considered for each biological process

Genome duplication: synthesis of deoxyribonucleotides, DNA unwinding, synthesis and ligation of Okasaki fragments and sliding clamp assembly.

Transcription: synthesis and polymerization of ribonucleotides, transcript length and half-life (rRNA, VSG, mRNA and SL RNA), nucleosome displacement, splicing.

Proteome maintenance: amino acid polymerization, protein half-life and degradation Membrane doubling: synthesis of phospholipids and ergosterol

Synthesis of sugar nucleotides: average glucidic composition

Synthesis of polyphosphates: synthesis of short-chain and long-chain polyP, polyP translocation

Transmembrane transport: transport of ions and amino acids

Cell motility: flagellar beating, dynein sliding displacement and power stroke Activation and recruitment of vesicles: rate of vesicle formation, Rab assembly Maintenance of mitochondrial membrane potential: Fo-ATPase activity

Information about some of the processes listed here is very complete. However, for some other processes in the trypanosome major gaps exists in our knowledge, while for still other ones very little information is available. Where possible, quantitative information was taken from other organisms, or assumptions have been made. Where this has been done, it is mentioned in the text and tables.

**Table 13.** Databases used in this work.

| Database | Web address | Reference |
| --- | --- | --- |
| TriTrypDB | https://tritrypdb.org/tritrypdb/app | (Aslett et al., 2010) |
| Bionumbers | https://bionumbers.hms.harvard.edu/search.aspx | (Milo et al., 2010) |

## Acknowledgements

This work was supported by: Fundação de Amparo à Pesquisa do Estado de São Paulo (FAPESP) grants 2017/16553-0 and 2021/12938-0 (awarded to AMS), Conselho Nacional de Pesquisas Científicas e Tecnológicas (CNPq) grant 307487/2021-0 (awarded to AMS) and Wellcome Trust grant 222986/Z/21/Z (awarded to JFN and AMS). ROOS, MBA, SM, AMM, FSD, RBMMG, LM, LLM and RWA are FAPESP fellows.

## Author Contributions

JFN, ROOS, MBA, SM, AMM, FSD, RBMMG, LM, LLM, RWA, PAMM and AMS compiled, organized and analyzed the data, designed and produced all tables and figures, and wrote essential parts of this manuscript. PAMM and AMS conceived the topic, scope and general organization of the manuscript. JFN, JHR, PAMM and AMS contributed to the revision and edition of the manuscript. PAMM and AMS made the final revision and edition. All authors approved the submitted version.

